# Novel Molecular Classification of Muscle-Invasive Bladder Cancer Opens New Treatment Opportunities

**DOI:** 10.1101/327114

**Authors:** Lucía Trilla-Fuertes, Angelo Gámez-Pozo, Guillermo Prado-Vázquez, Andrea Zapater-Moros, Mariana Díaz-Almirón, Jorge M Arevalillo, María Ferrer-Gómez, Hilario Navarro, Paloma Maín, Enrique Espinosa, Álvaro Pinto, Juan Ángel Fresno Vara

**Affiliations:** Biomedica Molecular Medicine SL, Madrid, Spain; Molecular Oncology & Pathology Lab, Institute of Medical and Molecular Genetics-INGEMM, Hospital Universitario La Paz-ldiPAZ, Madrid, Spain; Biostatistics Unit, Hospital Universitario La Paz-ldiPAZ, Madrid, Spain; Department of Statistics, Operational Research and Numerical Analysis, Universidad Nacional de Educación a Distancia (UNED).; Department of Statistics and Operations Research, Faculty of Mathematics, Complutense University of Madrid, Madrid, Spain.; Servicio de Oncología Médica, Hospital Universitario La Paz-ldiPAZ, Madrid, Spain; Biomedical Research Networking Center on Oncology-CIBERONC, ISCIII, Madrid, Spain

**Keywords:** computational analyses, immune status, molecular subtypes, muscle-invasive bladder cancer, personalized medicine

## Abstract

**Background:** Muscle-invasive bladder tumors are associated with high risk of relapse and metastasis even after neoadjuvant chemotherapy and radical cystectomy. Therefore, further therapeutic options are needed and molecular characterization of the disease may help to identify new targets.

**Objective:** The aim of this work is to characterize muscle-invasive bladder tumors at molecular levels using computational analyses.

**Design, Settings and Participants:** The TCGA cohort of muscle-invasive bladder cancer patients was used to describe these tumors.

**Outcome Measurements and Statistical Analysis:** Probabilistic graphical models, layer analyses based on sparse k-means coupled with Consensus Cluster, and Flux Balance Analysis were applied to characterize muscle-invasive bladder tumors at functional level.

**Results:** Luminal and Basal groups were identified, and an immune molecular layer with independent value was also described. Luminal tumors had decreased activity in the nodes of epidermis development and extracellular matrix, and increased activity in the node of steroid metabolism leading to a higher expression of androgen receptor.

This fact points to androgen receptor as a therapeutic target in this group. Basal tumors were highly proliferative according to Flux Balance Analysis, which make these tumors good candidates for neoadjuvant chemotherapy. Immune-high group had higher expression of immune biomarkers, suggesting that this group may benefit from immune therapy.

**Conclusions:** Our approach, based on layer analyses, established a Luminal group candidate for androgen receptor inhibitor therapy, a proliferative Basal group which seems to be a good candidate for chemotherapy, and an immune-high group candidate for immunotherapy.

**Patient Summary:** Muscle-invasive bladder cancer has a poor prognosis in spite of appropriate therapy. Therefore, it is still necessary to characterize these tumors to propose new therapeutic targets. In this work we used computational analyses to characterize these tumors and propose treatments.

## Introduction

Bladder cancer is estimated to account for 81,190 new cases and 17,240 deaths in the United States in 2018 [1]. Muscle invasive bladder cancer (MIBC) is characterized by a high risk of relapse and metastasis [2], The standard treatment consists of neoadjuvant chemotherapy followed by radical cystectomy. Nevertheless, neoadjuvant chemotherapy is a cisplatin-based schedule that is associated with significant toxicity.

Some patients do not have benefit form this approach, with tumors progressing despite the administration of chemotherapy. Therefore, these patients will be receiving a toxic and unnecessary treatment, as well as delaying a potentially curative treatment, such as surgery. Unfortunately, we do not have reliable biomarkers to guide us in patient selection for these therapies. Several translational studies have aimed to identify subgroups of patients with different clinical behavior.

Choi *et al.* identified three groups of MIBC (luminal, basal and p53-like) with different response to neoadjuvant chemotherapy [3]. The Cancer Genome Atlas (TCGA) developed a molecular classification of MIBC based on RNAseq data and hierarchical cluster analysis [4], In this work, five different groups were established: luminal-papillary (which included luminal tumors with papillary histology), luminal-infiltrated (characterized for lymphocyte infiltration), luminal, basal/squamous (also with lymphocyte infiltration) and a small neuronal group.

Seiler *et al.* associated TCGA molecular subtypes with response to neoadjuvant chemotherapy in a new cohort of patients [5]. Basal tumors appeared to benefit most from neoadjuvant chemotherapy, whereas luminal immune infiltrated tumors had a worse prognosis. However, these findings are not compelling enough to drive clinical decisions, so further insight into the molecular biology of MIBC is needed.

Data was analyzed using three mathematical methods that have proved to be very useful in other fields. Probabilistic graphical models (PGM) can identify differences in biological process among tumors [6–9]. Mathematical classification methods, such as sparse k-means [10] and Consensus Cluster [11], previously demonstrated their utility in the establishment of tumor subtypes [6]. On the other hand, Flux Balance Analysis (FBA) is a widely used approach for modeling biochemical networks. FBA could be used to calculate tumor growth rate [12].

In this study, data from the TCGA cohort were analyzed through PGM and computational analysis to characterize MIBC at the functional level.

## Material and methods

### TCGA cohort: data pre-processing

TCGA RNAseq data from patients with MIBC and treated with surgery alone were used to perform this study. Patients treated with neoadjuvant therapy were initially excluded for computational analysis. For survival analysis (which relied on clinical information), subjects who had received targeted therapies or radiotherapy were excluded, as well as those with M1 disease or missing T-stage information.

Log2 of the data was calculated and genes appearing in less than 75% of the samples were discarded. Missing values were imputed using a normal distribution with Perseus software [13], as previously described [7].

### Probabilistic graphical models

2,345 genes with highest variation in expression (standard deviation >2) were selected to build the PGM. The PGM method is compatible with high-throughput data with correlation as associative coefficient, as previously described [6–9]. Briefly, gene expression data was used without other *a priori* information and the analyses were done using *grapHD* package [14] and R v3.2.5 [15]. The resulting network was split into several branches and the most representative function of each branch was established by gene ontology analyses using DAVID webtool, as previously described. [16]. Functional node activities were calculated by the mean expression of the genes related to the main function assigned to each node.

### Biological layer analyses

Sparse k-means [10] and Consensus Cluster Analysis (CCA) [11] were used to explore molecular groups in the TCGA MIBC data, as previously described [6]. Sparse k-means assigns a weight to each variable, based on its relevance in the sample classification. Then, a CCA using variables that were selected by the sparse k-means method was applied to define the optimum number of groups for each case. The sparse k-means-CCA workflow was performed several times to explore the presence of independent informative molecular layers. Once relevant genes for one molecular layer were identified, these genes were removed from the dataset and the sparse k-means-CCA workflow was performed again, allowing the identification of different layers of molecular information and establishing different classifications based on various molecular features. Gene ontology analyses were performed for each layer to derive functional information.

### Flux Balance Analysis and flux activities

FBA was used to build a computational model that predicts tumor growth rates. COBRA Toolboox available for MATLAB [17], and the human whole metabolic reconstruction Recon2 [18] were used. The “biomass” reaction included in the Recon2 was designated as objective function. As described in previous works [8,9], expression data was included into the model by solving GPR rules and using a modified E-flux algorithm [9,19].

As in previous works [9], flux activities for each pathway were calculated by the sum of fluxes for all reactions involved in one pathway as defined in the Recon2. Then, comparisons between luminal and basal groups were performed using a Mann-Whitney test.

### Statistical analyses

GraphPad Prism v6 was used for statistical analyses. All p-values were two-sided and considered statistically significant under 0.05. Network analyses were done using Cytoscape software [20].

## Results

### Data pre-processing and patient selection

The TCGA cohort included 427 patients. Ten patients who had received neoadjuvant chemotherapy were excluded, leaving 417 subjects for subsequent analyses. Patients treated with targeted therapy or radiotherapy; M1 at diagnosis or no specified muscle-invasive type in the database were excluded from the analyses involving clinical data. Therefore, 178 patients were used for survival analyses (Sup Figure 1).

### Patients and samples characteristics

Data from 178 patients included in the clinical analyses are summarized in Sup Table 1. Median of overall survival, considering a five years period of follow-up, is 1,270 days and there were 73 death events during this time.

### Functional network

PGM were used to build a network, as previously described [6–8, 21] and the resulting network was functionally characterized. The network included 13 branches or functional nodes, one of them without a main relevant biological function and one with two different main biological functions: cytochrome metabolism and steroid metabolism (Figure 1).

**Figure 1:**
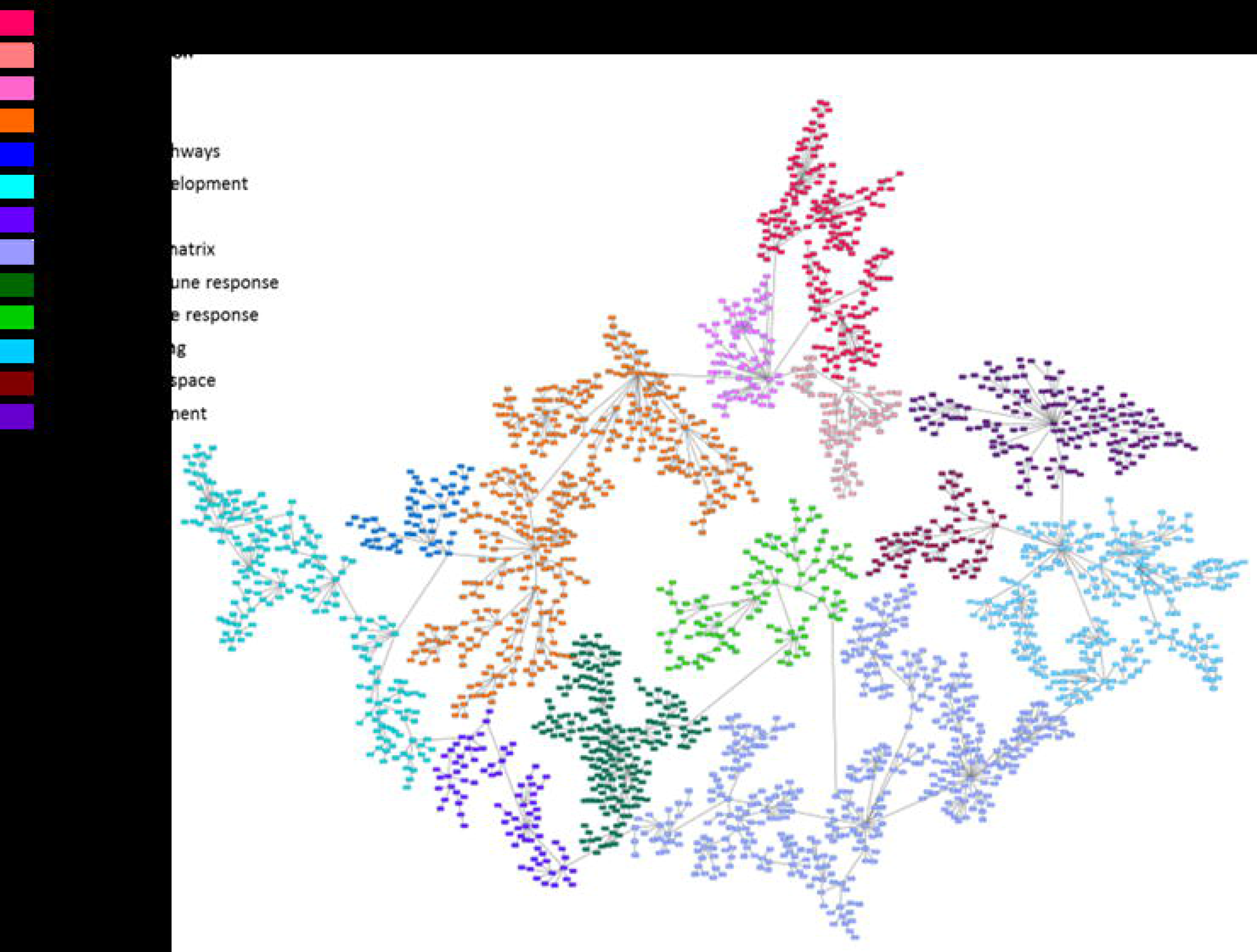
Probabilistic graphical model graph showing the network functional structure. Each node is named as its gene ontology main function identified.

### Biological layer analyses

The sparse k-means-CCA workflow defined 16 different layers of information (Sup Table 2). The first three layers had different ontologies and were further analyzed. Layers 4 to 13 had similar ontologies that one of the first three layers, so they were dismissed.

### Layer 1: Extracellular exosome and epidermis development

Layer 1 was based on 75 genes, which were mainly related with extracellular exosome, epidermis development and sodium ion homeostasis. This layer divided patients into two different groups. Group 1.1 included 260 patients (62.35%) and was characterized by lower expression of genes included in the epidermis development and extracellular matrix nodes. Group 1.2 included 157 patients (37.64%) and showed higher expression of genes included in the epidermis development and extracellular matrix nodes (Sup Figure 2). The TCGA classification of MIBC establishes the existence of both luminal and basal groups. Interestingly, our Group 1.1 had a higher expression of KRT20, GATA3 and FOXA1 genes, all of them luminal biomarkers, whereas Group 1.2 had a higher expression of KRT5, KRT6 and KRT14 genes, all of them basal biomarkers (Figure 2 and 3). So, from now on, Group 1.1 will be called Luminal group and Group 1.2, Basal group. Luminal tumors had a trend towards better survival than basal tumors, although the difference was not statistically significant (p= 0.1210, HR= 0.6969) (Sup Figure 3).

**Figure 2:**
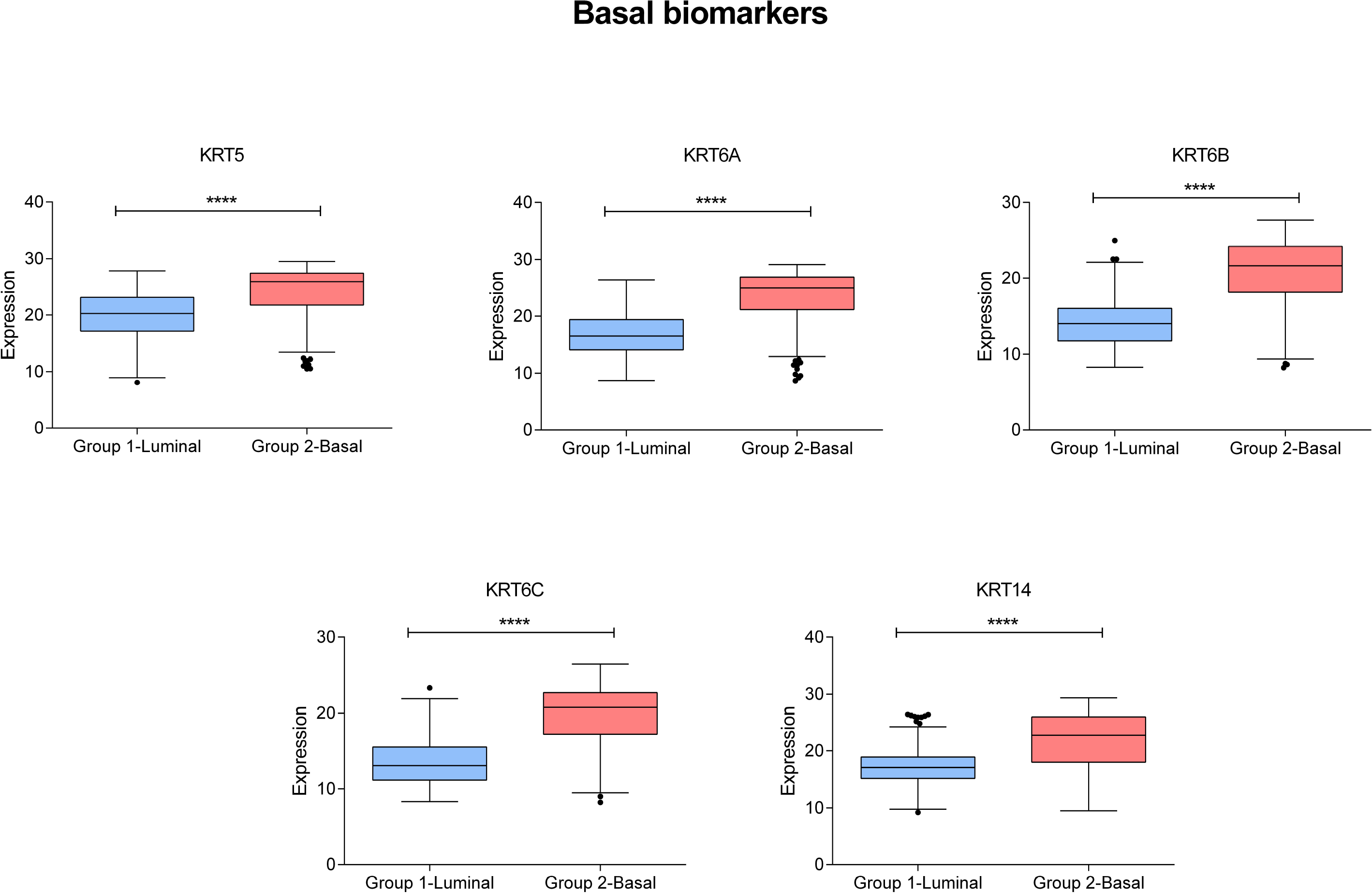
Expression of Basal biomarkers in Group 1.1 and Group 1.2.

**Figure 3:**
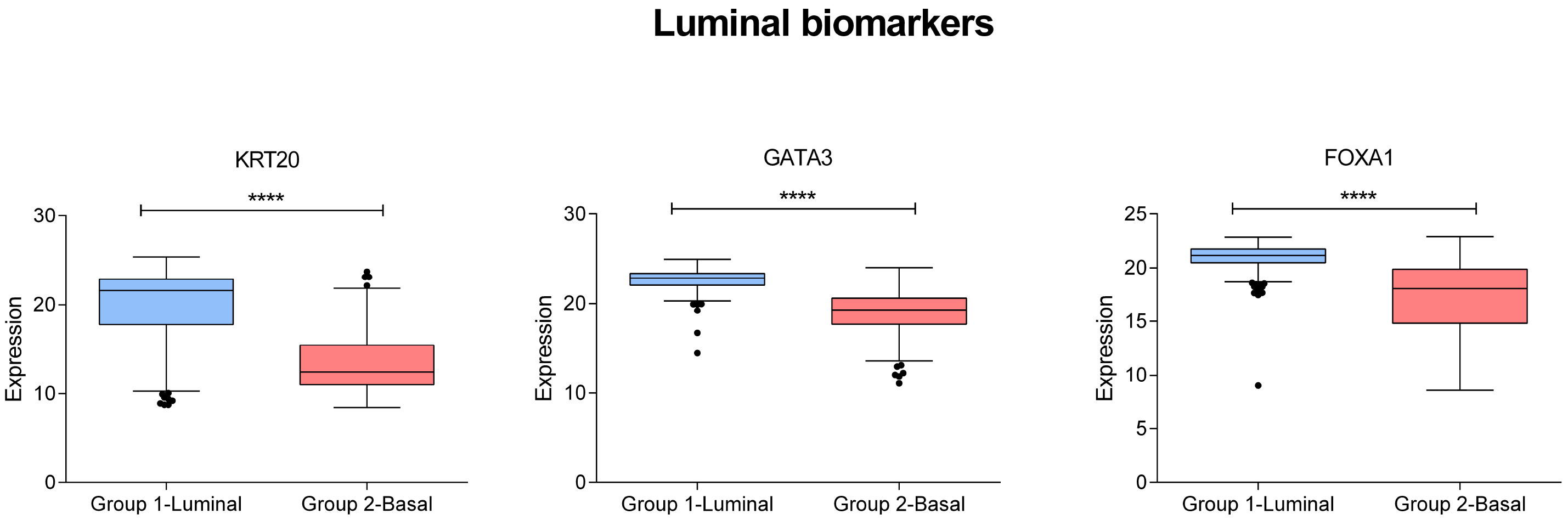
Expression of Luminal biomarkers in Group 1.1 and Group 1.2

Functional node activity for each node was calculated and compared between these two groups as previously described [6–8]. There were significant differences between luminal and basal subgroups in cytochrome metabolism, steroid metabolism, membrane, DNA binding, stem cell pathways, epidermis development, growth, extracellular matrix, adaptive immune response, innate immune response, extracellular space, and CNS development functional node activity (Sup Figure 4).

Differences in steroid metabolism node between luminal and basal tumors led us to evaluate the expression of androgen receptor (AR) in both groups. Interestingly, Luminal tumors showed higher expression of the AR gene (p<0.0001, fold change = 2.669) (Figure 4).

**Figure 4:**
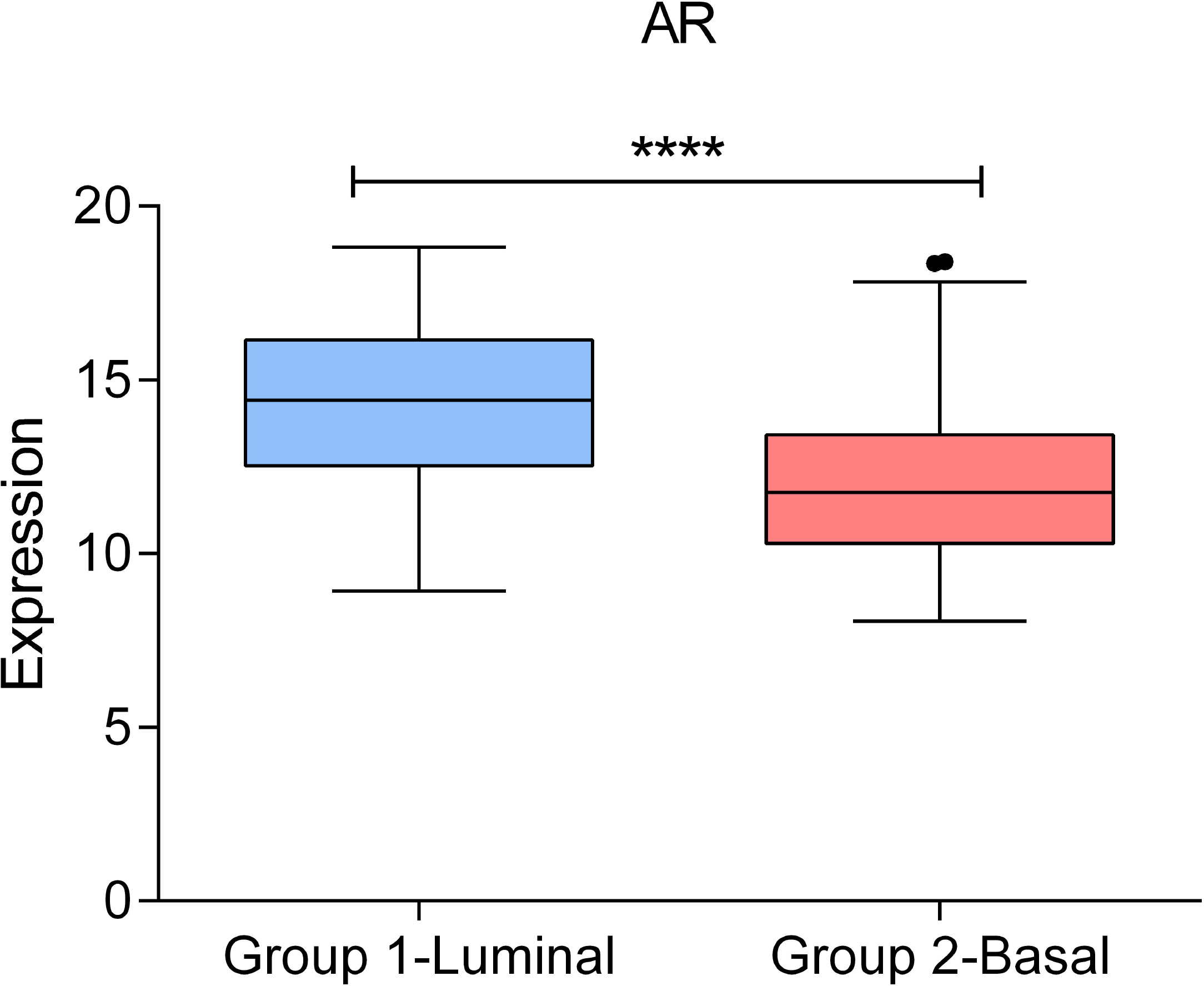
Androgen receptor expression in Luminal and Basal group.

### Layer 2: Extracellular space

Layer 2 was based on 82 genes mainly related with extracellular space. Layer 2 classifieds 268 patients (64.3%) in Group 2.1 and 149 patients (35.7%) in Group 2.2. Group 2.1 was characterized by higher expression of the extracellular matrix node and lower expression of the cytochrome metabolism node. Group 2.2 had the opposite expression pattern (Sup Figure 5). Group 2.1 had better prognosis than Group 2.2(Sup Figure 6).

### Layer 3: Immune

Layer 3 was based on 66 genes mainly related with inflammatory immune response. This layer divided patients into two groups. Group 3.1 included 215 patients (51.55%) and Group 3.2, 202 patients (48.44%). Group 3.1 was characterized by a high expression of innate and adaptive immune response nodes so, from now on, it will be called immune-high group. Group 3.2 was characterized by a low expression of innate and adaptive immune response nodes and will be called immune-low group (Sup Figure 7). The TCGA study used CD274 and CTLA4 to define immune infiltration in both luminal and basal tumors. In our new groups, these two immune biomarkers were more expressed in the immune-high group (Figure 5). In addition, the immune-low group had better prognosis, although the difference was not statistically significant (Sup Figure 8). As expected, the node activities of immune nodes were higher in the immune-high group (Sup Figure 9). Both the basal and the luminal groups contained immune-high and immune-low tumors, although the basal group had less immune-high tumors (Sup Figure 10).

**Figure 5:**
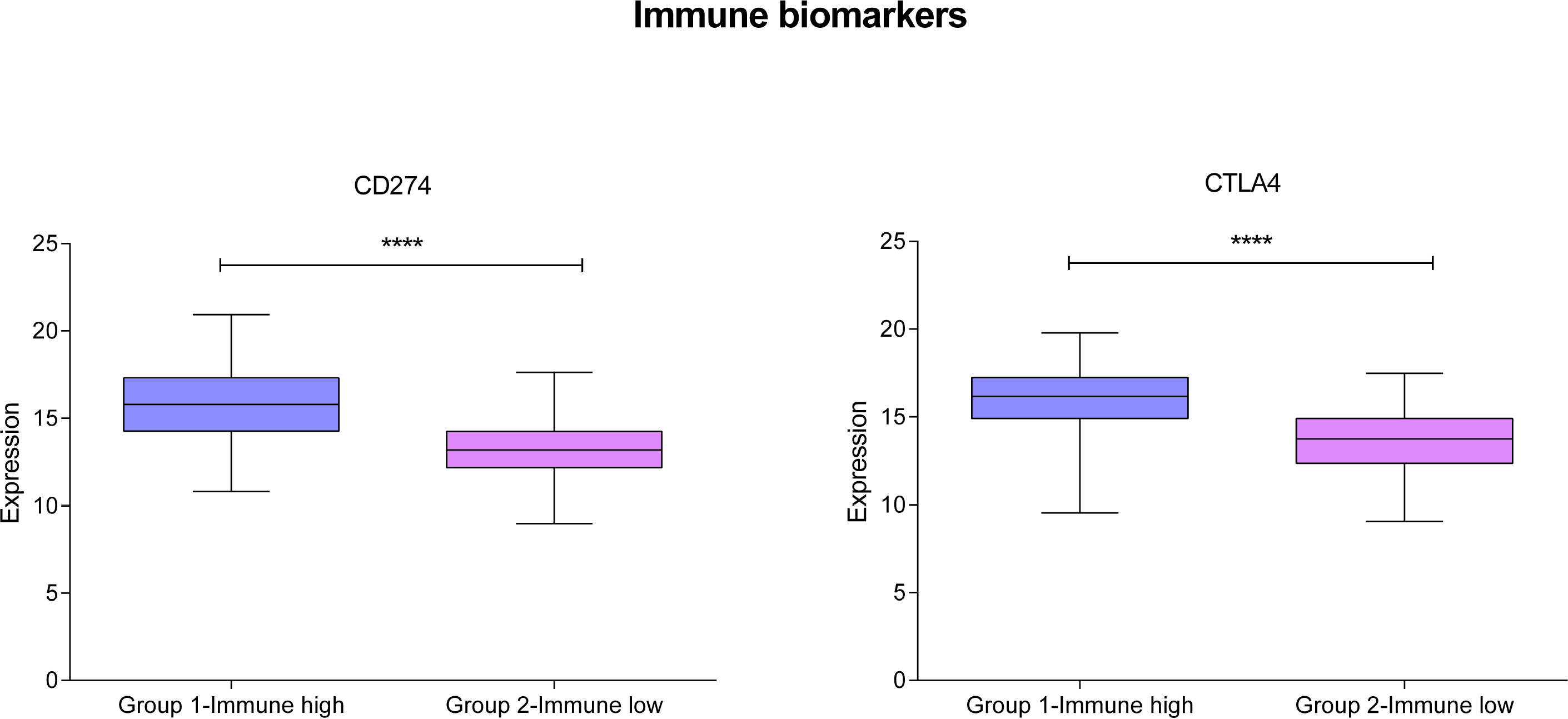
Expression of immune biomarkers in our immune groups.

### Layers 14 and 15

The first three layers provided distinct ontology information, but layers 4 to 13 contained redundant information. Layer 14 (translation) and layer 15 (chemical synapsis) provided no additional grouping information (Sup Figure 11).

### Flux Balance Analysis growth predictions and flux activities

FBA was used to study tumor growth and compare it between the layer-defined groups. According FBA predictions, basal tumors were more proliferative than luminal tumors (Figure 6) (p<0.0001).

**Figure 6:**
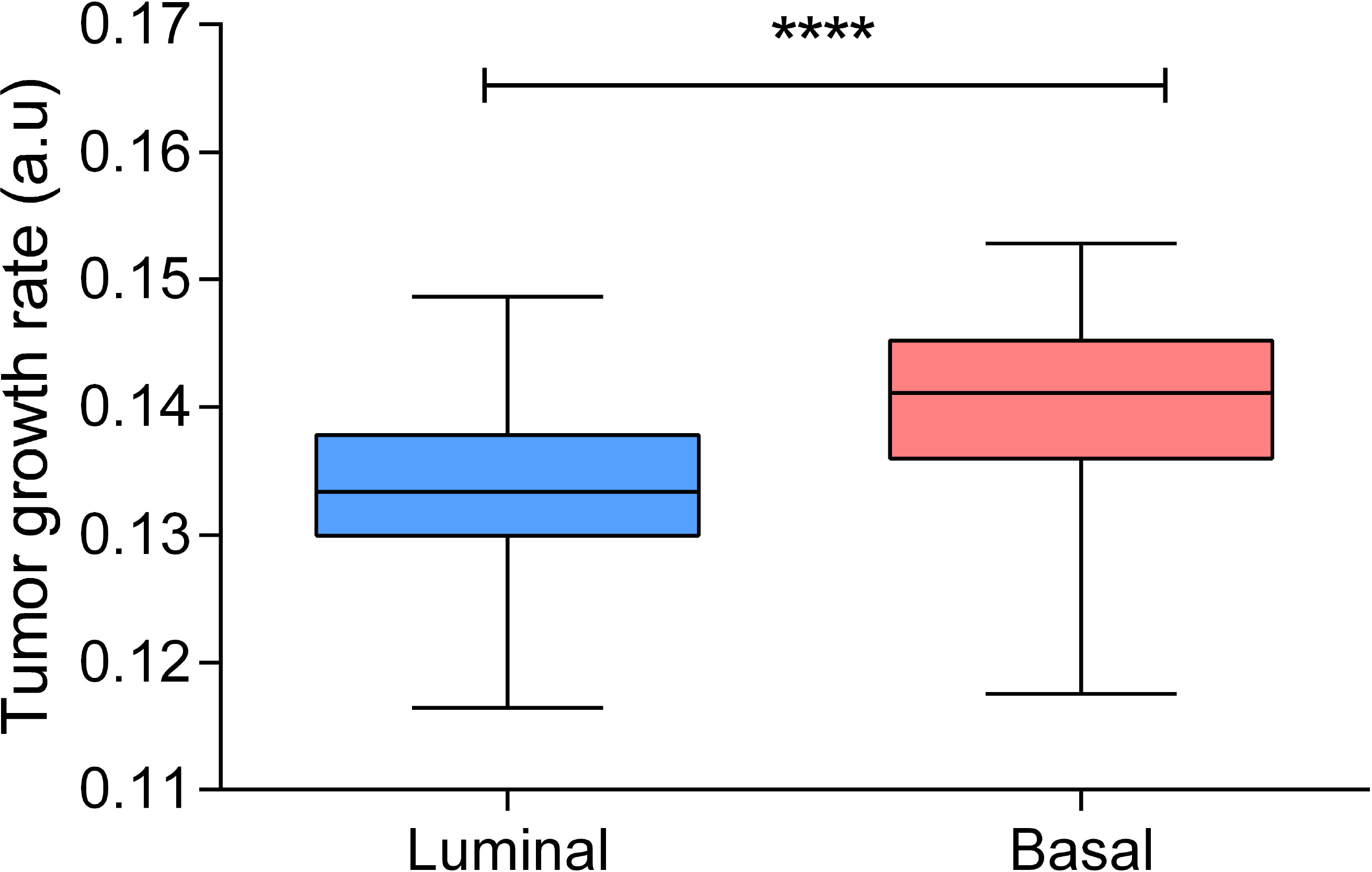
Tumor growth rate predicted by FBA in luminal and basal tumors.

Flux activities were calculated for each metabolic pathway and compared between basal and luminal tumors to determine differential pathways as described previously [21], Luminal tumors had a higher androgen and steroid metabolism flux activity. Differences were also detected in aminosugar metabolism, coA synthesis, galactose metabolism, glycolysis, hyaluronan acid metabolism, lysine, methionine, NAD, nucleotide savage, oxidative phosphorylation, phosphatidyl inositol, pyrimidine synthesis, R group, triacylglycerol and vitamin B6 metabolism (Sup Figure 12)

### Comparison with TCGA classification

The TCGA classification mixes histological, immune and luminal-basal information, establishing three luminal groups: luminal, luminal-papillary, and luminal-infiltrated (by lymphocytes), one basal group characterized by an immune positive status, and a small group called neuronal [4], Our classification established the immune information as an independent layer divided between luminal and basal groups, i.e., both of them had immune-high and immune-low tumors. Therefore, not all basal tumors were classified as immune positive. Additionally, the TCGA luminal-papillary group, which is defined solely by histological features, had immune-high tumors when the layer classification was applied. Percentages were similar between both classifications, although we did not identify a neuronal group (Sup Table 3).

## Discussion

MIBC has a poor prognosis, with over 50% of relapses in spite of appropriate therapy [22], Therefore, it is still necessary to characterize MIBC in order to propose new therapeutic targets. With the aim of characterize MIBC patients at functional and molecular level, PGM, layer analyses and FBA were performed in this study to provide insight into the molecular features of MIBC.

The PGM unveiled the functional structure of tumors, which has been previously described by our group [7–9]. This allows the study of gene expression data from biological and functional points of view. As an example of consistency in this regard, cytochrome P450 and UDP-glucuronosyltransferase were related to androgen receptor in the same node, and it is known that androgens modulate the expression of these enzymes [23].

Layer analyses provided different information about molecular features of the tumors, as for example, about the cellular adhesion process and about the immune status. The first layer divided MIBC tumors into Luminal and Basal. Luminal tumors presented a higher steroid metabolism node activity. Indeed, AR gene presents higher expression in Luminal tumors, suggesting the utility of AR as a possible therapeutic target. AR was previously related with bladder cancer progression [24] and *in vitro* studies showed that a siRNA against AR decreased proliferation of AR-positive bladder cancer cell lines but had no effect on AR-negative cells [25]. Therefore, patients with Luminal MIBC tumors, characterized by high expression of AR, could be candidates for therapy with AR inhibitors.

Luminal tumors had a higher flux activity of androgen and steroid metabolism pathway, which agrees with the results found in node activity. Luminal tumors also had a higher flux activity at glycolysis pathway so they may respond to drugs targeting metabolism as metformin, which has been shown to reduce growth in bladder cancer cells [26].

On the other hand, FBA predicted that, as it has been seen in breast cancer [8], basal tumors are more proliferative than luminal tumors. It is established that basal breast cancer tumors have a good response to chemotherapy because they are more proliferative [27, 28]. Based on the FBA results, the previous knowledge in basal breast tumors, and taking into account that this cohort is chemotherapy-naïve, basal patients may be good responders to chemotherapy as it was previously suggested by Seiler *et al.* [5]. Proliferation has been determined in other tumor types through gene expression, but data in bladder carcinoma are scarce. FBA allows not only to study proliferation but also to compare metabolic pathways.

The third layer identified an immune high expression group with high expression of CTLA4 and CD274, which may be a group of patients candidates of receiving immunotherapy, given that CD274, also known as PD-L1, and CTLA4 are the basis of current immunotherapy [29]. Interestingly, immune high tumors had a worse prognosis than immune low tumors, according with Seiler *et al.* results, which suggested that luminal immune infiltrated tumors had a worse prognosis [5].

Percentage distribution between luminal and basal tumors was comparable in the TCGA classification and the layer classification. However, the TCGA classification mixes immune, histological and molecular information. Our approach, on the contrary, establishes two independent informative layers: molecular and immune; and it rendered some interesting findings that complement the TCGA classification: for instance, 10% of basal tumors had an immune-low status, whereas in the TCGA classification all basal tumors had an immune-positive status. With the arrival of immunotherapies to the clinic, it could be useful to characterize the immune status of tumors and establish groups with differential immune features to drive treatment decisions.

The study has some limitations. Our findings should be validated in an independent cohort. Publications including expression and clinical data are scarce, so validation would rather be performed in a prospective study. On the other hand, the existence of a neuronal group could not be confirmed. As neuronal group accounted for a minority of cases in the TCGA study, maybe we should have analyzed a larger population. Finally, although our results suggest that some drugs may work better in specific groups, this should be prospectively validated. Response to chemotherapy, for instance, has been related to multiple factors and the proliferation profile may not be enough to identify responders.

## Conclusions

Computational analyses found different levels of information in gene expression data from MIBC: one of these levels refers to immune features, whereas the other corresponds to the previous classification into luminal/basal subgroups. Our classification may have therapeutic implications for the treatment of MIBC.

### Take Home Message

We used computational analyses in a muscle-invasive bladder cancer cohort and we defined independent molecular and immune features in these tumors that allow us to suggest therapeutic targets.

## Acknowledgments

This study was supported by Instituto de Salud Carlos III, Spanish Economy and Competitiveness Ministry, Spain and co-funded by the FEDER program, “Una forma de hacer Europa” (PI15/01310). LT-F is supported by Spanish Economy and Competitiveness Ministry (DI-15-07614). GP-V is supported by Conserjería de Educación, Juventud y Deporte of Comunidad de Madrid (IND2017/BMD7783). The funders had no role in the study design, data collection and analysis, decision to publish or preparation of the manuscript.

## Disclosures

JAFV, EE and AG-P are shareholders in Biomedica Molecular Medicine SL. LT-F and GP-V are employees of Biomedica Molecular Medicine SL. The other authors declare that they have no competing interests.

## Supplementary table legends

Sup Table 1: Clinical patients’ characteristics.

Sup Table 2: Main gene ontology term defined for each sixteen layers obtained by the sparse k-means-CCA workflow.

Sup Table 3: Percentages of patients assigned to each group in TCGA and layer classification.

### Supplementary figure legends

Sup Figure 1: Flowchart for patient selection.

Sup Figure 2: PGM’s graph heatmap showing differences between Group 1.1 (Luminal) and Group 1.2 (Basal). Green= underexpressed, Red= overexpressed.

Sup Figure 3: Kaplan Meier analysis comparing Luminal and Basal MIBC tumors clinical evolution.

Sup Figure 4: Functional node activities comparison between Luminal and Basal group.

Sup Figure 5: PGM’s graph heatmap showing differences between Group 2.1 and Group 2.2. Green= underexpressed, Red= overexpressed.

Sup Figure 6: Kaplan Meier analysis comparing Group 2.1 and Group 2.2 clinical evolution.

Sup Figure 7: Heatmap showing differences between Group 3.1 (immune-high) and Group 3.2 (immune-low). Green= underexpressed, Red= overexpressed.

Sup Figure 8: Survival curves obtained for immune groups.

Sup Figure 9: Immune node activities in immune groups.

Sup Figure 10: Concordance between classification of layer 1, which divided tumors into Luminal and Basal, and layer 3, which divided tumors into immune-high and immune-low group.

Sup Figure 11: Heatmap showing differences between groups defined in layers 14 and 15. Green= underexpressed, Red= overexpressed.

Sup Figure 12: Flux activities of luminal and basal groups.

